# Integrin β1–Talin1 at focal adhesions underpin uncontrolled endothelial cell enlargement in live cerebral cavernous malformation vasculature

**DOI:** 10.1101/2025.11.25.688491

**Authors:** Mikaela S. Keyser, Teodor E. Yordanov, Jason A. da Silva, Tevin CY. Chau, Elysse K. Morris, Scott Paterson, Neil I. Bower, Katarzyna Koltowska, Gregory J. Baillie, Cas Simons, Robert G. Parton, Kelly A. Smith, Benjamin M. Hogan, Anne K. Lagendijk

## Abstract

Cerebral cavernous malformations (CCMs) are vascular anomalies caused by loss of CCM gene function and consequent hyperactivation of MEKK3-KLF2/4 signaling in endothelial cells. Excess integrin β1 activity has been associated with lesion growth, yet the precise mechanistic and biomechanical roles of the integrin-KLF2/4 hierarchy in CCM cellular pathogenesis are not fully understood. Using live imaging of endothelial Vinculin in Ccm1-deficient zebrafish, we demonstrate excessive, mechanically active focal adhesions in an *in vivo* model of CCM pathology. We validate this in CCM1-deficient endothelial cells and show a redistribution of mechanical tension from cell-cell junctions to focal adhesions. Genetic deletion of Talin1 to decouple focal adhesions from the cell cortex and inhibit integrin β1 signaling demonstrates the integrin β1-Talin1 complex is essential for vascular malformations in *ccm1* mutants by driving endothelial cell enlargement. We show integrin β1-Talin1 act independent or downstream of KLF2/4, rather than upstream as previously suggested. Thus, we reposition the role of integrin β1-Talin1 in CCM pathogenesis, demonstrating an essential function in driving cell enlargement, which underpins lesion growth.

## Introduction

Cerebral cavernous malformations (CCMs or cavernomas) are vascular lesions that primarily develop in low flow venous capillaries of the central nervous system (CNS)^1,2^. Lesions can arise sporadically or be a consequence of inherited mutations in one of three genes that encode CCM proteins: CCM1 (KRIT1), CCM2 (OSM) or CCM3 (PDCD10)^3–6^. Biochemical studies have identified that CCM proteins can interact to form the CCM signaling complex (CSC)^7–10^. Endothelial restricted loss of the individual CCM proteins in mice leads to similar vascular malformations^11–13^, indicating that CCM proteins mainly function as part of the CSC. Progressed CCM lesions in patients are characterized as distinctively dilated vascular regions with a multi-lumen cavern morphology. The ECs that line such progressed lesions are enlarged and thin with compromised cell-cell adhesion^3,14^. Mechanistically, CCM proteins inhibit RhoA-ROCK activity, thereby controlling phosphorylation of myosin light chains (pMLC), acto-myosin contractility, and cell- cell junction integrity^13,15–17^. In *Ccm1/Krit1* and *Ccm2* heterozygous mice, vascular leakage phenotypes were effectively resolved by treatment with either simvastatin^17^, a Rho GTPase inhibitor, or Fasudil^18^, a ROCK inhibitor. Overall, these studies led to the general appreciation that increased contractility through RhoA-ROCK contributes to CCM cellular phenotypes that underpin fragility of lesions and enhance the risk of rupture and haemorrhaging^13,15,19^.

Discovery of the MEKK3–KLF2/4 signaling axis as a key transcriptional driver of CCM pathogenesis added a new layer of mechanistic understanding^20–23^. Reducing the genetic dosage of *Mekk3* or *Klf2/4* prevented lesion formation in *Ccm1/Krit1* mice^21^. To place this in context of established findings on uncontrolled RhoA–ROCK signaling, pMLC was shown to be reduced upon Klf2/4 loss, indicating that changes in RhoA–ROCK occur downstream of KLF2/4^21^. Since CCM1-CCM2 associate and stabilize the integrin β1 inhibitor (Integrin cytoplasmic domain-associated protein 1, also referred to as integrin subunit β1 binding protein 1, or ITGB1BP1)^24,25^, and functional studies in cultured cells demonstrated that enhanced integrin β1 activity can induce RhoA and cell shape changes^26,27^, integrin β1 gained interest as a major mechanical player in CCM. In accordance with *in vitro* data, lesion burden of *Krit1^flox/flox^;Pdg@-iCreERT2* mice, whereby *Krit1/Ccm1* is knocked out in the endothelium, was significantly reduced upon treatment with 9EG7, an integrin β1 inhibitory antibody^28^. Similarly, in zebrafish, knocking down *integrin β1b* with morpholinos partially rescued cardiac morphogenesis in *ccm2* mutants^22^. These studies also probed the effect of integrin β1 inhibition on KLF2/4 activity, however with conflicting results, concluding that inhibiting integrin β1 *in vitro* either restored^22^ or had no effect^28^ on excessive KLF2/4 mRNA in CCM. These discrepancies underscored the need to clarify the molecular hierarchy between integrin β1 and the KLF2/4 in CCM pathogenesis.

Integrin β1 is essential for cell–matrix adhesion in a multitude of cell and tissue types, including blood vessels^29–31^. Intracellularly, integrin receptors are bound by talin and kindlin adaptor proteins that are required for integrin signaling and mechanical coupling to the actin cytoskeleton^32,33^. ICAP1 competes with talin and kindlin adaptor proteins for integrin β1 binding^34,35^. When CCM1 is lost, ICAP1 is destabilized and undergoes proteasomal degradation^25,26^, which would increase accessibility for talins and kindlins to interact and activate integrins. Integrin complexes are found at focal adhesions (FA) and FA number, size and dynamics are commonly used *in vitro* as read-outs for integrin activity^33,36,37^. Live visualization of endothelial FAs *in vivo* has been challenging, which complicates direct investigation of KLF2/4 transcriptional activity in relation to FA dynamics in CCM vasculature. To address this limitation, we here employ high-resolution live imaging of endothelial Vinculin at FAs^38^ in Ccm1-deficient zebrafish and reveal an increase in FAs that temporally emerges after Klf2 induction. With *in vivo* genetic studies we define the integrin-KLF2/4 hierarchy and reveal key CCM cellular phenotypes that are regulated by integrin β1-Talin1.

## Results

### Live visualization of ectopic focal adhesions in *ccm1^uq6al^* mutants

To monitor morphological and transcriptional changes in endothelial cells devoid of Ccm1, we characterized a novel *ccm1^uq6al^*allele that was isolated from a previously conducted forward genetic screen^39^. This allele harbors a single nucleotide nonsense mutation at serine428 in the ARD domain, which leads to a premature stop (Fig. 1a). When examining overall morphology of *ccm1^uq6al^* mutant embryos, and found that homozygous mutants are indistinguishable from siblings at 24 hpf (Fig. 1b). However, by 48 hpf, *ccm1^uq6al^* mutants had developed a severe enlargement of the heart, which is characteristic to Ccm loss-of-function in embryonic zebrafish^22,40–48^ (Fig. 1b). We next utilized our previously validated *Tg(fli1ep:vinculinb-eGFP)^uq2al^* transgenic line^38^ to visualize FAs in the developing posterior cardinal vein (PCV) (Fig. 1c). This, to our knowledge, represents the first *in vivo* visualization of FAs in live CCM vasculature. We observed that FAs were not altered in *ccm1^uq6al^* mutants at 30 hpf (Fig. 1d, e). By 48 hpf, however, we identified large Vinculinb-eGFP clusters in *ccm1^uq6al^* mutant PCVs that were not observed in siblings (Fig. 1d, e). Vinculin is known to be recruited to both FAs and adherens (cell-cell) junctions (AJs) to assists in maintaining actin connectivity under increased acto-myosin tension^49–51^. AJs were challenging to distinguish in the PCV, particularly in *ccm1^uq6al^* mutants. To faithfully uncouple Vinculinb-eGFP at FAs versus AJs, we examined Vinculinb-eGFP expression in the common cardinal vein (CCV) (Fig. 1c), where AJs are more pronounced. Live imaging of the CCV at 48 hpf uncovered a similar increase in Vinculinb-eGFP positive FAs in *ccm1^uq6al^* mutants (Fig. 1f, g). Collectively this data therefore provides the first *in vivo* evidence that Ccm1 loss leads to an increase in endothelial FAs. The presence of Vinculin further indicates that these FAs are under mechanical tension.

**Figure 1:**
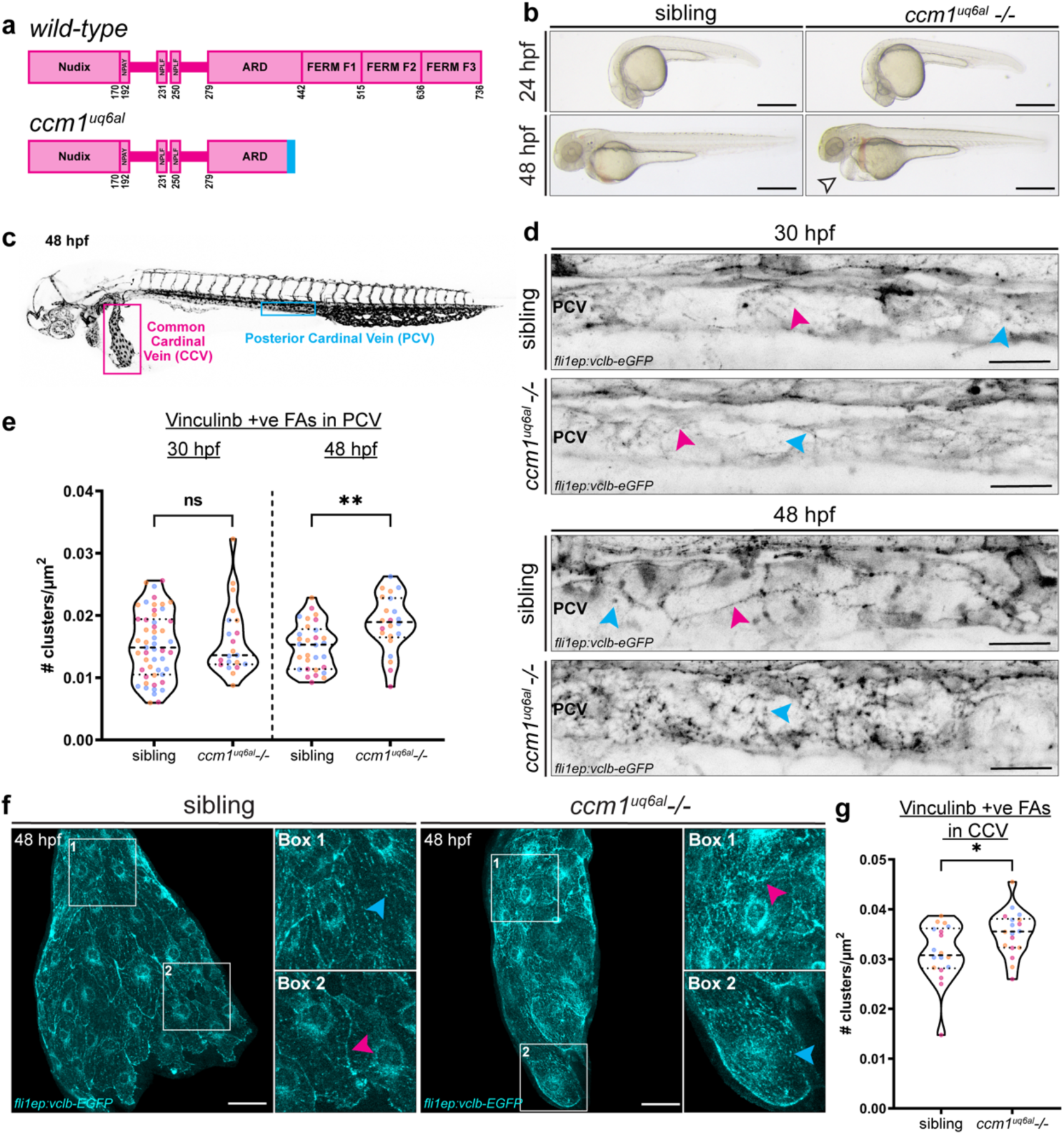
Characterization of *ccm1^uq6al^* mutants demonstrated increased focal adhesion formation *in vivo*. **(a)** Schematic representation of wild-type CCM1 protein and truncation in *ccm1^uq6al^* mutants. **(b)** Sibling and *ccm1^uq6al^ -/-* embryos at 24 hpf were phenotypically indistinguishable, however at 48 hpf *ccm1^uq6al^ -/-* embryos developed cardiac oedema (arrowhead) and lack blood circulation (scale bar = 500 μm). **(c)** Blood vessel network (grey) in a 48 hpf *Tg(kdrl:eGFP)* zebrafish embryo, highlighting the CCV (magenta box) and PCV (blue box). **(d)** Vinculinb-eGFP protein localization in the PCV at 30 hpf (top) and 48 hpf (bottom). Blue arrowheads indicate Vinculin at FAs and magenta arrowhead indicates Vinculin at AJs (scale bar = 30 μm). **(e)** Quantification of the number of Vinculin-eGFP positive FA clusters in the PCV at 30 hpf and 48 hpf, showing an increase in PCV clusters at 48 hpf; 3 biological replicates, n(sibling, 30 hpf) = 51, n(*ccm1-/-,* 30 hpf) = 21, n(sibling, 48 hpf) = 30, n(*ccm1-/-*, 48 hpf) = 23. Unpaired two-tailed Student’s t-test. **(f)** Vinculinb-eGFP protein localization in the CCV at 48 hpf. Blue arrowheads indicates Vinculin at FAs and magenta arrowhead indicates Vinculin at AJs (scale bar = 40 μm). **(g)** Quantification of the number of Vinculin-eGFP positive FA clusters in the CCV at 48 hpf; 3 biological replicates, n(sibling) = 18, n(*ccm1-/-*) = 18. Unpaired two-tailed Student’s t-test. Data represented using violin plots, small circles indicate individual data points for each replicate (color matched to replicate). ns=no significant difference, *p=0.0332, **p=0.0021.

### Selective recruitment of vinculin to focal adhesions over adherens junctions in CCM1-deficient endothelial cells

We investigated these observations *in vitro* in CCM1 LOF HUVECs, in which CCM1 protein is lost upon CRISPR-Cas9 targeting^52^ (Fig. 2a). We validated that ITGB1 protein is increased in CCM1 LOF monolayers (Fig. 2b, c). To determine the impact on FA formation and maturation, we visualized pPaxillin and Vinculin by immunostaining. We identified an increase in the number of pPaxillin-positive FAs in CCM1 LOF HUVECs (Fig. 2d, e), and these FAs were significantly larger, as reported previously^26,27,52–54^ (Fig. 2d, f). This increase in FA size and the presence of pPaxillin suggest that these are mature FAs that are under greater tension, in accordance with our *in vivo* data (Fig. 1d-g). To explore this further, we assessed Vinculin expression at FAs and AJs, as an indicator of adhesion sites that are under high tension. Quantitative analysis of the ratiometric abundance of Vinculin relative to VE-cadherin at AJs and to pPaxillin at FAs uncovered a mechanical shift with increased Vinculin recruitment to these enlarged FAs and a significant decrease in Vinculin at AJs (Fig. 2d, g, h). Since CCM proteins are known to inhibit Rho- ROCK activity and the formation of actin stress fibers^15,19^, our *in vivo* and *in vitro* analysis indicates that this mechanical imbalance might be a consequence of enhanced contractility of these FA-coupled stress fibers after loss of CCM1.

**Figure 2:**
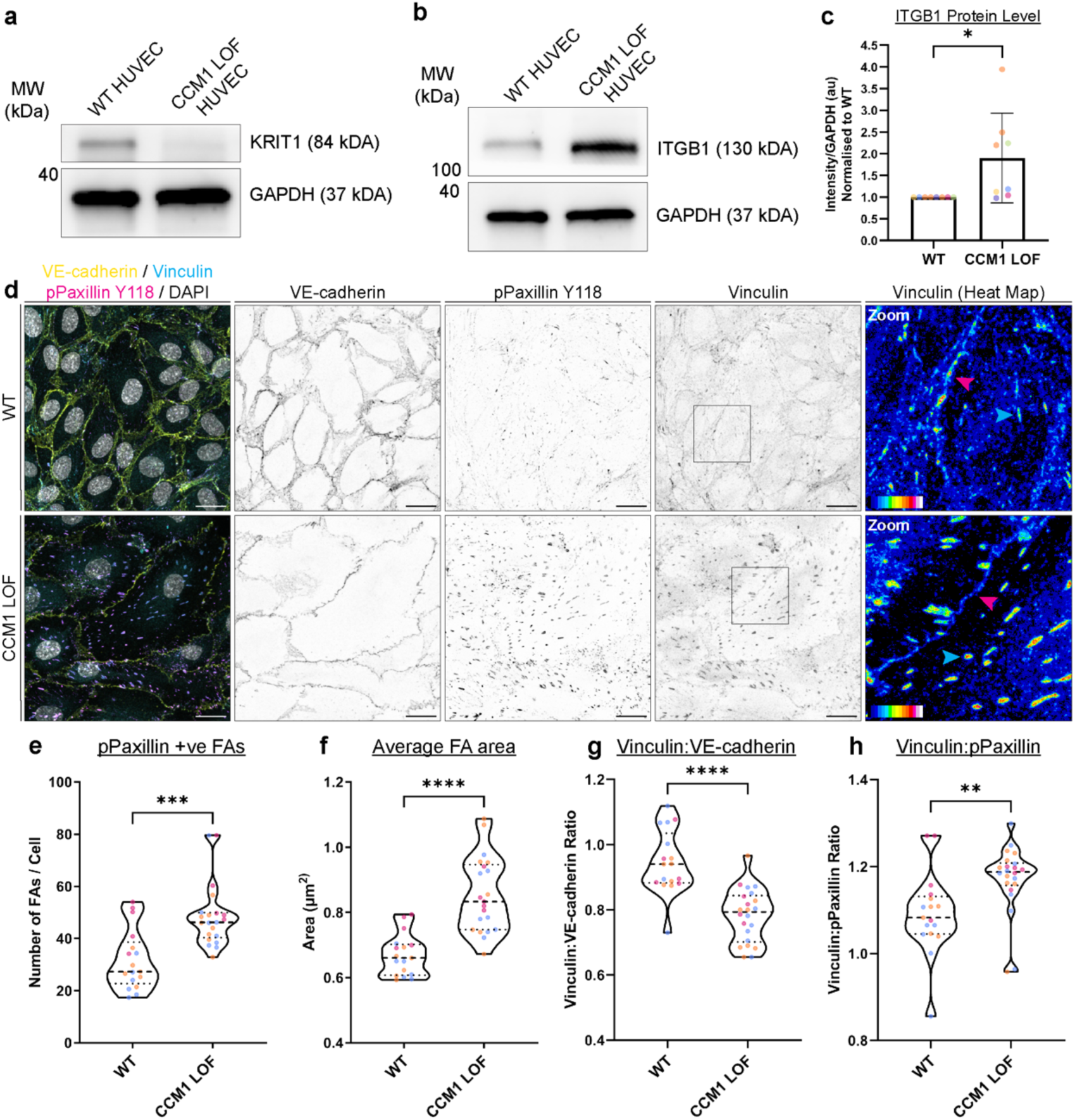
CCM1 LOF ECs have excess Vinculin positive focal adhesions. **(a)** Western blot of WT and CCM1 LOF ECs demonstrated a loss of CCM1 (KRIT1) protein expression. **(b)** Western blot of WT and CCM1 LOF HUVECs demonstrated an increase in Integrin β1 protein expression. **(c)** Quantification of Integrin β1 protein expression, corrected to loading control and normalized to WT HUVECs. 6 biological replicates, unpaired two-tailed Student’s t-test. **(d)** Maximum intensity projection of confluent monolayers of WT and CCM1 LOF ECs with immunofluorescent staining phospho-Paxillin (Tyr118, magenta), Vinculin (cyan), VE-cadherin (yellow), and nuclei (DAPI, grey). Heatmap of Vinculin expression, with magenta arrowhead indicating Vinculin at AJs and cyan arrowhead indicating Vinculin at FAs (scale bar = 20 μm). **(e)** Quantification of the number of FAs per EC and **(f)** average FA area comparing WT ECs to CCM1 LOF ECs, based on pPaxillin expression. **(g)** Quantification of the ratio of Vinculin over VE-cadherin, indicating cadherin AJs under mechanical tension **(h)** Quantification of the ratio of Vinculin over pPaxillin, indicating FAs under mechanical tension. 3 biological replicates, 5-8 images per replicate, with each data point representing an average of 4-24 cells per image. Unpaired two-tailed Student’s t-test (F, G) and unpaired two-tailed Student’s t-test with Welch’s correction (E, H). Data represented using violin plots or bar plots, small circles indicate individual data points for each replicate (color matched to replicate). *p=0.0332, **p=0.0021, ***p=0.0002, ****p<0.0001.

### Loss of Talin1 reduces cardiovascular phenotypes in *ccm1^uq6al^*mutants

To interrogate how these mature and mechanically active FAs may contribute to CCM defects *in vivo,* we combined our *ccm1^uq6al^*mutant with a *tln1^uq1al^* (*talin1*) mutant allele^38^. Talin proteins are required to connect integrins to the actin cytoskeleton and transduce chemical and mechanical signaling^32,55,56^. In mice, Talin1 loss in CCM1/*Krit1-*deficient endothelium caused extensive cerebral hemorrhaging which precluded cellular analysis of lesion endothelium^28^. We have previously reported that endothelial FAs fail to form in our zebrafish *tln1^uq1al^* strain, yet overall cardiovascular defects are not as severe as those observed in Talin1 knockout mice^57–59^ and thus this model allows us to visualize the cellular consequences of compromised integrin signaling in live Ccm1-deficient animals. Major phenotypic hallmarks for Ccm loss in zebrafish include the formation of enlarged blood vessels, cardiac ballooning, and a failure of the heart to undergo cardiac looping and form a defined atrio-ventricular canal (AVC)^22,23,42,44–46,48^. We examined these cardiovascular phenotypes in progeny of *ccm1^uq6al^;tln1^uq1al^* double heterozygotes (double mutants will be further referred to as *ccm1*;*tln1*). As reported previously, at 2 dpf, *tln1^uq1al^* mutants could be distinguished by cranial hemorrhaging and a smaller heart, accompanied by cardiac edema^38^ (Fig. 3a, b). In *ccm1;tln1* double mutant embryos we noted a reduction in cardiac ballooning compared with homozygous *ccm1^uq6al^* mutant embryos (Fig. 3a, b). Although *ccm1;tln1* mutant hearts were smaller, atrial and ventricular morphology remained compromised and the AVC did not form (Fig. 3b). These morphological defects impacted function of *ccm1;tln1* hearts, leading to a complete lack of blood flow in *ccm1;tln1* double mutants, as observed in *ccm1^uq6al^* mutants. This lack of perfusion likely explains why we did not observe cranial hemorrhaging in *ccm1*;*tln1* mutants, in contrast to *tln1^uq1al^*mutants where the cranial vasculature remains perfused at this developmental stage. To determine the impact of Talin1 loss on vascular phenotypes, we next performed live imaging of the lateral dorsal aorta (LDA). LDA width is a common read-out in the field for CCM related vessel dilation^22,44^. As reported previously for other Ccm mutant strains^22,44^, the LDA in *ccm1^uq6al^*mutants was significantly dilated, with an increase in LDA vessel width and vessel area (Fig. 3c-e). In *ccm1;tln1* double mutants, the LDA did not expand and instead these vessels were thinner and smaller than wild-type LDAs (Fig. 3c-e). Together, this analysis of cardiovascular morphogenesis in *ccm1;tln1* double mutants uncovered that Talin1, and by extension cell-matrix adhesion, is required for cardiac and vascular dilation when Ccm1 is lost.

**Figure 3:**
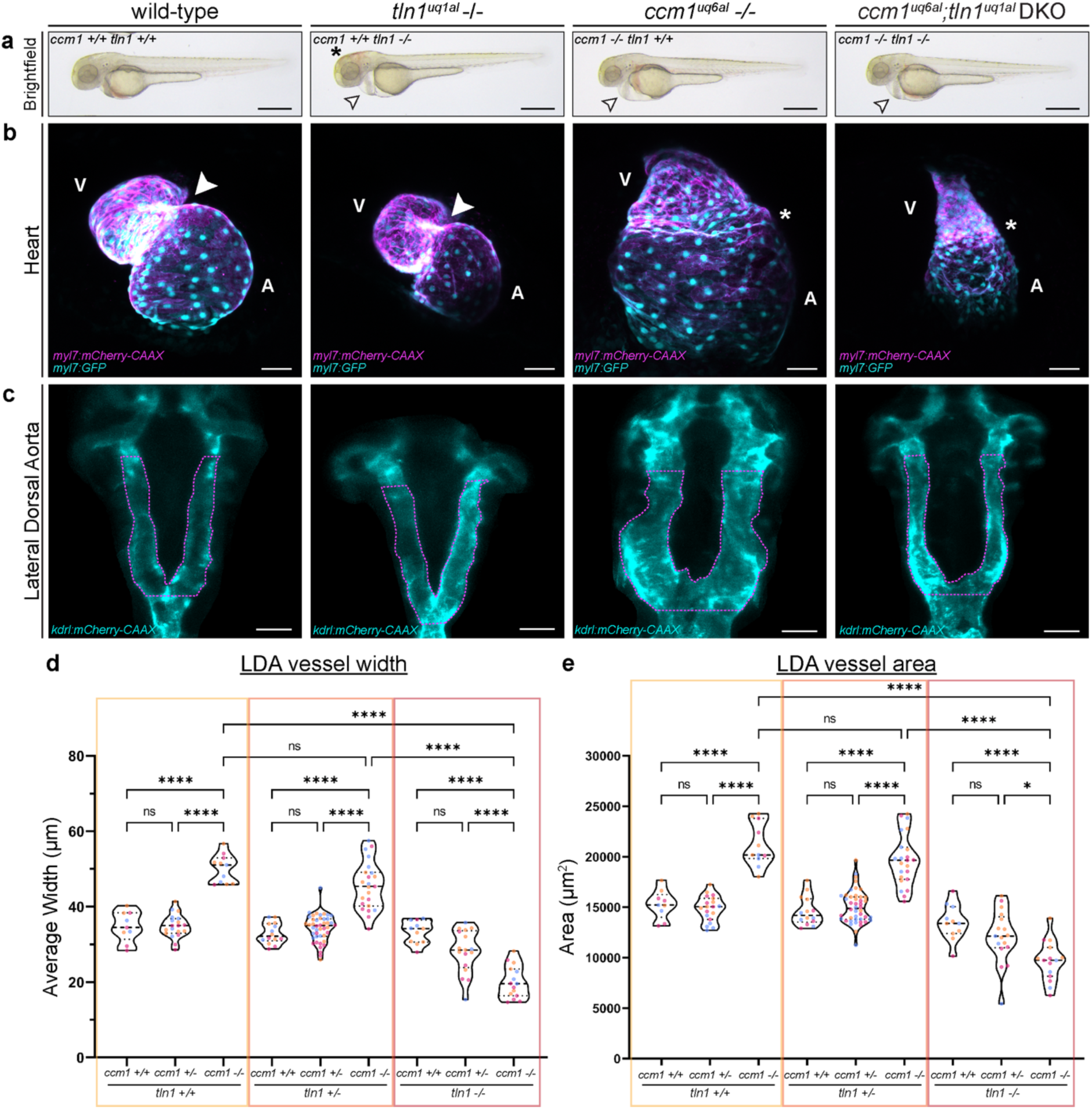
Loss of Talin1 recovers cardiovascular abnormalities in *ccm1^uq6al^* -/- embryos. **(a)** Brightfield images showing gross morphology of wild-type, *tln1^uq1al^ -/-*, *ccm1^uq6al^ -/-*, and *ccm1; tln1* double mutants at 2 dpf, with arrowheads indicating cardiac oedema and asterisk indicating brain hemorrhaging (scale bar = 500 μm). **(b)** Maximum intensity projections of zebrafish embryos hearts at 2 dpf, showing that loss of Talin1 inhibits cardiac ballooning observed in *ccm1^uq6al^* -/- embryos (scale bar = 50 μm). A = atrium, V = ventricle, white arrowhead = AVC, asterisks indicate AVC absence. **(c)** Maximum intensity projections of the LDA at 2 dpf, demonstrating significant vessel dilation in *ccm1^uq6al^-/-*, which does not occur in *ccm1; tln1* double mutants (scale bar = 50 μm and LDA area is outlined in magenta). **(d)** Quantification of LDA average vessel width and **(e)** average LDA vessel area; 3 biological replicates, n(*ccm1+/+;tln1+/+*) = 9, n(*ccm1+/+;tln1+/-*) = 14, n(*ccm1+/+;tln1-/-*) = 11, n(*ccm1+/-;tln1+/+*) = 19, n(*ccm1+/-;tln1+/-*) = 44, n(*ccm1+/-;tln1-/-*) = 17, n(*ccm1-/-;tln1+/+*) = 11, n(*ccm1-/-;tln1+/-*) = 23, n(*ccm1-/-;tln1-/-*) = 15. One- way ANOVA with Tukey’s multiple-comparisons test. Data represented using violin plots, small circles indicate individual data points for each replicate (color matched to replicate). ns=no significant difference, ***p=0.0002, ****p<0.0001.

### Integrin activation does not drive pathological KLF2/4 activity in CCM

Knockdown of *integrin β1b* in zebrafish *ccm2* mutants has previously been shown to partially rescue cardiac morphogenesis^22^, similar to what we observe in our *ccm1;tln1* genetic model. Mechanistic data indicated that integrin β1 acted upstream of Klf2 in CCM, inducing KLF2/4 mRNA levels in CCM^22^. Since Talin1 is essential for integrin β1 signaling^32,33,56^, we hypothesized that Talin1 would be essential to drive this integrin β1-Klf2/4 cascade in CCMs. In zebrafish, *klf2a* and *klf2b* are the functional orthologues of KLF2 and KLF4 in mammals^60,61^. We generated a *klf2a* reporter transgene, *Tg(klf2a:H2B-eGFP)^uq5al^*, analogous to previously published *klf2a* reporter line that faithfully reports pathological *klf2a* induction in zebrafish^62^. We first examined *klf2a* expression in the PCV of *ccm1^uq6al^* mutants by live imaging at 24 and 48 hpf. Quantitative analysis of nuclear *klf2a* expression showed that *klf2a* reporter activity was significantly upregulated in *ccm1^uq6al^* mutants compared with siblings at 24 hpf (Fig. 4a, b). Notably, this is prior to the onset of blood flow, the formation of excess FAs (Fig. 1d, e), and distinguishable cardiovascular defects (Fig. 1b), underscoring the primary role for Klf2 in CCM. By 48 hpf, despite a complete lack of blood flow in *ccm1^uq6al^* mutants, *klf2a* levels remained increased in *ccm1^uq6al^*mutants, as reported previously by others at this time point^22,44^ (Fig. 4a, b). To determine whether loss of Talin1 impacts *klf2a* induction in *ccm1^uq6al^* mutants, we next quantified *klf2a* levels at 2 dpf across all genotypes from *ccm1;tln1* double heterozygote progeny. In wild-type embryos, we observed baseline *klf2a* levels (Fig. 4c, d) whilst in *tln1^uq1al^* mutants *klf2a* reporter activity was significantly reduced (Fig. 4c, d). This is consistent with our previous finding that blood flow velocity is reduced in *tln1^uq1al^*mutants at 2 dpf^38^. Interestingly, loss of Talin1 did not prevent induction of *klf2a* in *ccm1^uq6al^*mutants (Fig. 4c, d). These differences in *klf2a* levels across genotypes were specific to the endothelium since *klf2a* was unchanged in muscle cells located in close proximity to the vasculature (Supplementary Fig. 1a, b). Together this *in vivo* analysis of *klf2a* reporter activity indicates that integrin β1-Talin1 function is not required to initiate Klf2a induction in CCM. Complementary *in vitro* analysis validated these findings. We used lentiviral delivery of an ITGB1 shRNA to knockdown integrin β1 (ITGB1) in combination with the CRISPR-Cas9 CCM1 lentivirus^52^, yielding double LOF ECs. We validated knockdown efficiency of the ITGB1 shRNA by immunofluorescence staining using an antibody that specifically labels active ITGB1, which showed a significant reduction in active ITGB1 in shRNA ITGB1 exposed ECs (Supplementary Fig. 2a, b). As observed in *ccm1^uq6al^* mutant embryos, in CCM1 LOF ECs, endogenous levels of nuclear KLF4 protein were significantly increased (Fig. 4e, f). Knocking down ITGB1 did not inhibit this KLF4 induction in CCM1 LOF ECs (Fig. 4e, f). This *in vitro* work therefore supports our *in vivo* findings and together the data demonstrates that pathological induction of KLF2/4 in CCM disease does not require integrin β1-Talin1.

**Figure 4:**
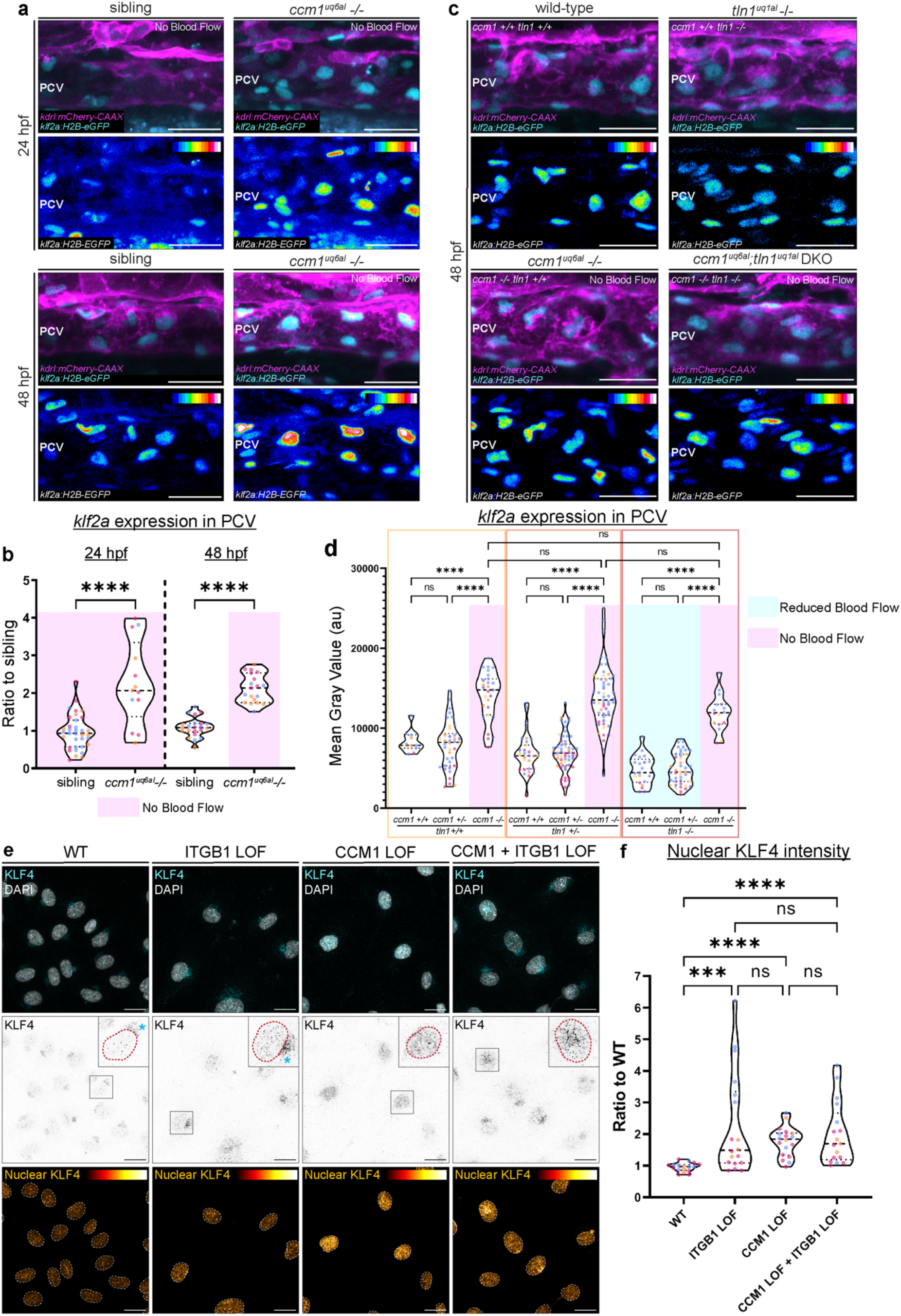
Integrin β1-Talin1 activation does not induce KLF2/4 in CCM *in vivo* and *in vitro*. **(a)** Maximum intensity projections of *klf2a* expression in the PCV at 24 hpf (top) and 48 hpf (bottom) in sibling and *ccm1^uq6al^ -/-* embryos (scale bar = 30 μm). **(b)** Quantification of *klf2a* intensity in the PCV of sibling and *ccm1^uq6al^ -/-* embryos at 24 hpf and 48 hpf; 3 biological replicates, n(sibling, 24 hpf) = 35, n(*ccm1-/-,* 24 hpf) = 13, n(sibling, 48 hpf) = 22, n(*ccm1-/-*, 48 hpf) = 19. One-way ANOVA with Tukey’s multiple- comparisons test. Magenta box = no blood flow. **(c)** Maximum intensity projections of *klf2a* expression in the PCV of wild-type, *tln1^uq1al^ -/-,* ccm*1^uq6al^ -/-,* and *ccm1;tln1* double mutants at 48 hpf (scale bar = 30 μm). **(d)** Quantification of *klf2a* expression in the PCV at 48 hpf. Cyan box = genotypes with reduced blood flow in trunk vessels, magenta box = genotypes with no blood flow; 5 biological replicates, n(*ccm1+/+;tln1+/+*) = 11, n(*ccm1+/+;tln1+/-*) = 32, n(*ccm1+/+;tln1-/-*) = 21, n(*ccm1+/-;tln1+/+*) = 38, n(*ccm1+/-;tln1+/-*) = 59, n(*ccm1+/-;tln1-/-*) = 43, n(*ccm1-/-;tln1+/+*) = 27, n(*ccm1-/-;tln1+/-*) = 54, n(*ccm1-/-;tln1-/-*) = 17. One-way ANOVA with Tukey’s multiple-comparisons test. **(e)** Top: Maximum z- projections of confocal images of confluent monolayers of WT, ITGB1 LOF, CCM1 LOF, and CCM1 + ITGB1 LOF ECs, fixed and stained for KLF4 (cyan) and nuclei (DAPI, grey). Middle: KLF4 only (grey) with inlay showing the nucleus outlined in red and perinuclear KLF4 is indicated by blue asterisks. Bottom: Red hot LUT for heatmap of nuclear KLF4 expression (scale bar = 20 μm). **(f)** Quantification of nuclear KLF4 intensity in WT, ITGB1 LOF, CCM1 LOF, and CCM1 + ITGB1 LOF ECs; 3 biological replicates, 6 images per replicate, with each data point representing 3-20 nuclei per image. One-way ANOVA with Tukey’s multiple-comparisons test. Data represented using violin plots, small circles indicate individual data points for each replicate (color matched to replicate). ns=no significant difference, *p=0.0332, **p=0.0021, ***p=0.0002, ****p<0.0001.

### Integrin β1 and Talin1 are required to drive EC enlargement in the absence of CCM1

Since integrin signaling is well known to drive cell spreading^33,37,63,64^, we hypothesized that functional integrin β1-Talin1 adhesions are required for excessive EC enlargement, a major hallmark of CCM loss^52,65–69^. To monitor EC size *in vivo*, we implemented our VE-cadherin transgenic line^70^, *TgBAC(ve-cad:ve-cad-TS)^uq17bh^*, and performed live imaging of VE-cadherin protein at endothelial cell-cell junctions. We examined the CCV at 48 hpf, since we identified that FAs are induced in *ccm1^uq6al^* mutant CCVs at this stage (Fig. 1f, g) and we can faithfully capture VE-cadherin positive cell-cell junctions across all genotypes. As expected, quantification of EC size revealed that ECs in the CCV of *ccm1^uq6al^* mutants were significantly larger (Fig. 5a, b). Whilst loss of Talin1 alone did not alter EC size, depleting Talin1 in *ccm1^uq6al^*mutants resulted in a recovery of EC size (Fig. 5a, b). We validated EC size rescue *in vitro* in CCM1;ITGB1 double LOF ECs. As previously reported, CCM1 LOF ECs were significantly larger (Fig. 5d, e). Interestingly, we also observed an increase in EC size upon ITGB1 knockdown, yet not to the extent of CCM1 LOF ECs (Fig. 5d, e). Knockdown of ITGB1 in CCM1 LOF ECs significantly reduced EC size back to that of ITGB1 LOF only ECs (Fig. 5d, e). We next determined whether cell size changes were associated with changes in VE-cadherin abundance at cell-cell junctions. Notably, loss of Talin1 resulted in increased VE-cadherin at AJs (Fig. 5c), suggesting uncoupling of focal adhesions might invoke strengthening of the AJs. In *ccm1;tln1* double mutants, however, VE-cadherin coverage was significantly reduced (Fig. 5c). This indicated that cell size rescue in *ccm1;tln1* double mutants is not a consequence of AJ stabilization but instead implicates that enlargement of CCM-deficient ECs is solely driven by excessive integrin adhesion and signaling at FAs.

**Figure 5:**
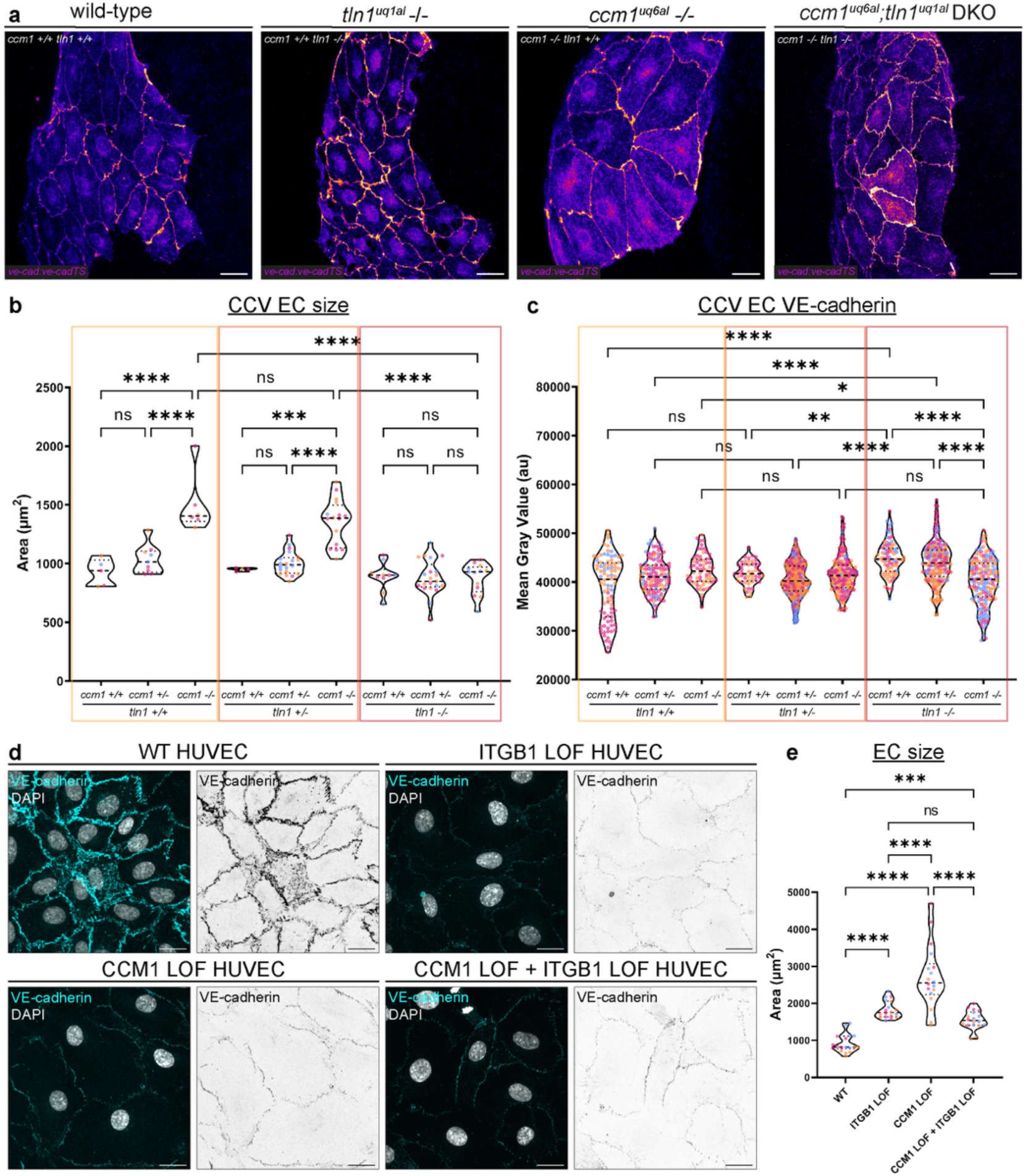
Integrin β1-Talin1 is required for EC size expansion of CCM1 deficient ECs *in vivo* and *in vitro*. **(a)** Maximum intensity projections of *TgBAC*(*ve-cad:ve-cadTS)* (VE-cadherin, fire) expression in the CCV in wild-type, *tln1^uq1al^ -/-*, *ccm1^uq6al^ -/-*, and *ccm1;tln1* double mutants at 2 dpf (scale bar = 25 μm). **(b)** Quantification of CCV EC size at 2 dpf; 3 biological replicates, n(*ccm1+/+;tln1+/+*) = 5, n(*ccm1+/+;tln1+/-*) = 4, n(*ccm1+/+;tln1-/-*) = 10, n(*ccm1+/-;tln1+/+*) = 9, n(*ccm1+/-;tln1+/-*) = 20, n(*ccm1+/-;tln1-/-*) = 21, n(*ccm1-/-;tln1+/+*) = 6, n(*ccm1-/-;tln1+/-*) = 16, n(*ccm1-/-;tln1-/-*) = 13. One-way ANOVA with Tukey’s multiple-comparisons test. **(c)** Quantification of VE-cadherin expression of the ECs of the CCV at 2 dpf; 3 biological replicates, Kruskall-Wallis test with Dunn’s multiple comparisons test. **(d)** Maximum z- projections of confocal images of confluent monolayers of WT, ITGB1 LOF, CCM1 LOF, ITGB1 and CCM1 LOF ECs, fixed and stained for VE-cadherin (cyan) and nuclei (DAPI, grey) (scale bar = 20 μm). **(e)** Quantification of EC size of WT, ITGB1 LOF, CCM1 LOF, ITGB1 and CCM1 LOF ECs in D; 3 biological replicates, 6 images per replicate, with each data point representing 2-19 cells per image. One-way ANOVA with Tukey’s multiple-comparisons test. Data represented using violin plots, small circles indicate individual data points for each replicate (color matched to replicate). ns=no significant difference, **p=0.0021, ***p=0.0002, ****p<0.0001.

## Discussion

Here we have visualized key CCM cellular hallmarks *in vivo* by live monitoring and show that mechanically active integrin-Talin1 complexes at FAs are essential for increased cell size, but not KLF2/4 transcription, as central aspects of CCM pathogenesis.

First, we imaged FA dynamics in live CCM vasculature and provided the first *in vivo* evidence that Vinculin-positive FAs are enriched in Ccm1-deficient veins. We show that Vinculin enrichment at FAs is accompanied by a reduction of Vinculin at the AJs. Vinculin is a mechanosensitive protein and is recruited to both AJs and FAs to strengthen actin connectivity when these adhesion sites are under increased tension^49–51^. Loss of Vinculin *in vivo* leads to a hyperpermeable vasculature, indicating that Vinculin recruitment to AJs is essential for adhesion stability and blood vessel integrity^71^. Depleting Vinculin from AJs in CCM therefore would likely contribute to enhanced permeability of lesions and hemorrhage risk. The increased abundance of Vinculin at FAs further provides molecular evidence in accordance with the observation that CCM proteins are required to promote AJ stabilization by ROCK2 whilst inhibiting ROCK1 activation and FA formation^47^. Traction force microscopy of angiogenic sprouting in a 3D hydrogel model revealed that CCM2-deficient ECs exert increased force on the ECM, further validating that these FAs are mechanically over-active and therefore would require Vinculin to support integrin-actin coupling^72^.

Second, we combined genetic knockout models for *ccm1* and *tln1* and revealed that Talin1 is required for vessel expansion and cardiac ballooning phenotypes. Similar cardiac improvements were observed when knocking down *integrin β1b* in Ccm2-deficient zebrafish^22^. With the integrin β1-KLF2/4 hierarchy not fully resolved or monitored *in vivo*, we examined if Talin1 is required for Klf2a induction in CCM. In *ccm1* mutants, Talin1 loss did not alter pathological induction of *klf2a,* and in a complementary approach *in vitro*, endogenous KLF4 protein remained increased in CCM1;ITGB1 double LOF ECs. Similarly, in an analogous experiment whereby CCM1-deficient ECs were treated with the integrin β1 inhibitor 9EG7, Klf4 mRNA remained upregulated^28^. From our *in vivo* and *in vitro* data, we propose that integrin β1- Talin1 does not induce pathological KLF2/4 activity in CCM, and that phenotypic rescue of cardiac ballooning and vessel expansion is not a consequence of normalizing KLF2/4.

Quantifications of EC shape instead uncover that integrin β1-Talin1 drive EC enlargement in CCM. Together, with the presence of ectopic Vinculin-enriched FAs, our findings suggest that activation of integrin β1 at FAs would induce cytoskeletal remodeling that promotes expansion of the cell membrane over the integrin β1-engaged substrate. When Talin1 is lost, mechanical coupling and signaling of integrins is inhibited, eliminating the ability of CCM cells to spread. Depletion of Talin1 in *Ccm1*/*Krit1* mice did not show improvements of CCM lesions, but induced severe hemorrhaging throughout the cerebral vasculature^28^, which might have masked the cellular consequences of effective Cre recombination of both *Krit1* and *Tln1* in mice^73^. Furthermore, it is worth exploring whether there are species-specific differences in functional redundancy amongst Talin proteins that might explain this difference in Talin1 requirement.

Notably, normalization of cell size in *ccm1;tln1* double mutants does not involve restoration of VE- cadherin at cell–cell junctions; instead, VE-cadherin levels are further diminished, indicating that CCM EC enlargement is primarily regulated at the level of FAs. Since cell size increase will contribute significantly to lesion growth, our data provides a mechanistic explanation that supports pharmacological inhibition of integrin β1 to inhibit CCM lesion development, as pioneered using 9EG7 in mice^28^. Whether integrin β1 interference could also be effective in reducing cell size in progressed CCM lesions remains to be determined.

Furthermore, the relative contribution of inside-out versus outside-in signaling to activate integrins in CCM is not fully understood. Transcriptomic profiling of ECs isolated from the heart and lungs of EC-specific *Klf2/4* KO mice, placed “ECM receptor Interaction” and “Focal Adhesion” as top dysregulated KEGG pathways in both EC cell types^74^, suggesting KLF2/4 regulates the expression of cell-matrix signaling genes There is also increasing evidence of ECM changes downstream of KLF2/4, that can alter CCM phenotypes non-cell autonomously. KLF2/4 induces expression of ADAMTS4 which leads to enhanced proteolytic cleavage of the white matter extracellular protein Versican^75^. This releases biologically active Versikines which were proven to be essential in CCM disease progression^75^. In a three-dimensional micro-vessel model for CCM, changes in the Hyaluronic acid (HA) turnover were suggested to alter the abundance of high-molecular weight (HMW) versus low-molecular weight (LMW) HA in the vascular ECM^52^. This altered cell size of KLF4 induced CCM1 deficient ECs with LMW HA inhibiting EC enlargement and the formation of excess pPaxillin positive FAs^52^. Investigation of WT-CCM2 mosaic sprouts in 3D hydrogels showed enhanced ECM breakdown by CCM2 deficient tip cells, and transduction of hypercontractility by direct mechanical coupling^72^. Excess ECM degradation and CCM-WT mechanical coupling led to an increase in stress fibers and integrin β1 FAs in neighboring WT stalk cells^72^. This phenotypic switch of WT ECs was proposed to contribute to the incorporation of these cells into growing CCM lesions, as observed by lineage tracing studies in mice^68,69^. It is intriguing to speculate that KLF2/4 induced changes in ECM turnover and release of extracellular signaling molecules alter lesion growth by modifying integrin signaling. Further exploration of CCM-WT mosaic vasculature *in vivo* would inform the spatiotemporal relevance of mechanical coupling and non-cell autonomous signaling during lesion initiation and progression. With a recent genome wide association study for coronary artery disease revealing CCM signaling genes as a novel CAD associated gene hub^76^, identification of key cellular and extracellular players that are controlled by CCM will not only advance opportunities for CCM therapeutics but may also inform a broader role for CCM proteins in controlling vascular health in both veins and arteries.

## Methods

### Zebrafish lines and handling

All zebrafish experimentation was adhered to the guidelines of the animal ethics committee of the University of Queensland (Permit 2022/AE000091). Zebrafish transgenic lines utilized in this study were *Tg(myl7:mCherry-CAAX; myl7:GFP)*^77^*, Tg(kdrl:mCherry-CAAX)^y1^*^71,78^*, Tg(klf2a:H2B-eGFP)^uq5al^*, *Tg(fliep:vclb- eGFP)^uq2al,^* ^38^*, TgBAC(ve-cad:ve-cadTS)^uq11bh,^* ^70^. All embryos were obtained through natural paired matings and were incubated at 28°C in a dark-phase incubator. Embryos were maintained in 10 cm petri dishes containing 1X E3 media (5 mM NaCl, 0.17 mM KCl, 0.33 mM CaCl2, 0.33 mM MgSO4) at a maximum density of n=60. At 24 hpf, all embryos were changed to E3 media supplemented with 0.0003% phenylthiourea (PTU) to prevent pigmentation.

### Zebrafish genotyping

Zebrafish embryos were genotyped post-imaging by removing embryos from agarose and placing each embryo into an individual microtube for DNA extraction. DNA was isolated using embryo lysis buffer (10mM Tris-HCl, 1mM EDTA, 50 nM KCl, 0.3% Tween-20, 0.3% IGEPAL CA-630), supplemented with 0.5% proteinase K immediately before use. Samples were incubated for 2 hours at 55°C, followed by 95°C for 5 minutes, before storage at 4°C. *ccm1^uq6al^* genotyping: The *ccm1^uq6al^* mutant allele was isolated from a forward genetic screen^39^. The allele harbors a single nucleotide nonsense mutation, resulting in a premature stop at serine428. *ccm1^uq6al^* genotyping was carried out by PCR amplification of a 229 bp region flanking the mutation in the *ccm1* gene. PCR products were enzymatically digested using Sau3AI for 4 hours at 37°C. Three Sau3AI restriction sites are present in wild-types, and one is removed because of the mutation in *ccm1^uq6al^*. Sau3AI digested PCR products were separated on a 2% SB agarose gel by electrophoresis. Genotypes are distinguished based on band sizes, with homozygous wild type resulting in a 130, 52, 33 and 13 bp bands, homozygous mutant displaying 182, 33 and 14 bp bands, and heterozygotes displaying 182, 130, 52, 33 and 13 bp bands. Primer sequences for genotyping PCR: CCM1-forward 5’- CGATGCTGTAAAAGAATCCGTATG-3’, CCM1-reverse 5’-CTCATGCCCTCCATGATCTG-3’. *tln1^uq1al^* genotyping: The *tln1^uq1al^* allele was generated by CRISPR genome editing, harboring an in-frame deletion of 6 amino acids and one amino acid substitution in the F1 FERM domain, resulting in Talin1 protein loss^38^. *talin1^uq1al^* genotyping was carried out by PCR as described previously^38^. In brief, a 124 bp region of the *talin1* gene was amplified using primers flanking the mutation site. The PCR products were separated via gel electrophoresis on a 2% SB agarose gel. Genotypes were identified by band size, with heterozygotes displaying two bands at 107 and 124 bp, homozygous wild type displaying a single band at 124 bp, and homozygous mutants displaying a single band at 107 bp. Primer sequences for genotyping PCR: Talin1-forward, 5’- CGCTGATTGGGATTTTAATGTGTATTCAG-3’, Talin1-reverse, 5’- CTGGTCTGAGTAGAAGAACTTTCTCC-3’.

### Live imaging of zebrafish embryos

Embryos were sorted for fluorescent transgenic expression at 24 hpf and 48 hpf using a Leica M165 series stereo microscope. Zebrafish embryos were immobilized using tricaine (Sigma Aldrich E10521- 50G, 0.08mg/ml) at the desired timepoint, before being embedded in 1% low melting point agarose (Sigma Aldrich, A9414, 1% in E3) in 35mm glass bottom dishes (Matek, P35G-1.5-20-C). Embryos were embedded laterally for trunk vasculature and common cardinal vein imaging, dorsally for brain vascular imaging, and ventrally for heart imaging. Confocal z-stacks were acquired using a Zeiss Axiovert 200 Inverted LSM 710 Meta Confocal Scanner, using a 40X 1.1 NA LD C-Apochromat water objective. Heart and brain images were captured utilizing a 20X 0.8 NA Plan Apochromat dry objective. All images were acquired at 1024 x 1024 dimensions and 16-bit. Brightfield images of zebrafish embryos were captured using a Nikon SMZ1270i with Tucsen Michrome 6 camera. For consistency, all trunk images were captured in a specific section above the yolk extension.

### Antibodies and dyes

The following antibodies were used: rabbit α-KRIT1 (CCM1) (WB 1:1000; Abcam, ab196025), rabbit α- GAPDH (WB 1:5000; Cell Signalling, 2118), mouse α-VE-cadherin conjugated to Alexa Fluoro 647 (IF 1:500, Becton Dickinson, 561567), rabbit α-phosphorylated Paxillin (Y118) (IF 1:250; Cell Signalling, 69363), mouse α-Vinculin (1:250; Sigma Aldrich, V9131), rabbit α-KLF4 (IF 1:100; Cell Signalling, 4038), rat α-mouse CD29 (9EG7) (IF 1:200; Becton Dickson, 550531), rabbit α-ITGB1 (WB 1:1000; Sapphire Bioscience, STJ93731). Secondary antibodies conjugated with Alexa-488, Alexa-555 or Alexa-647 for immunofluorescence staining were obtained from Invitrogen and used at a 1:1000 dilution for immunofluorescence. For Western blotting, secondary antibodies, horseradish peroxidase (HRP) conjugated goat α-rabbit (WB 1:5000; Sigma Aldrich, A0545) was used.

### Lentivirus vector design and generation

Transient CCM1 loss of function by lentivirus was performed as per Yordanov and Keyser et al. (2024)^52^. In brief, a guide RNA sequence targeting CCM1 (5’-GTATTCCCGAGAATTGAGACTGG-3’) based on Zhang Lab Guide Design Tools (https://www.zlab.bio/resources) was selected and cloned into lentiCRISPRv2 vector (Addgene, 52961) after having underwent digestion with BsmBI, dephosphorylation with Antarctic phosphatase, and purification (New England BioLabs, T1020) as per Zhang Lab’s GeCKO LentiCRISPRv2 Target Guide Sequence Cloning Protocol. The gRNA was phosphorylated and annealed before undergoing ligation with the linearized lentiCRISPRv2 plasmid. Ligated plasmids were then transformed into dH5a competent cells and grown on LB agar plates supplemented with ampicillin. A negative control ligation (vector-only) was included in all ligation and transformation experiments. Colonies were grown overnight before being picked and grown in LB broth supplemented with ampicillin. Each culture underwent DNA purification using Favorgen Plasmid DNA Extraction Mini Kit (Favorgen, FAPDE001-1) and the gRNA insert was verified by sequencing. Predesigned ITGB1 gene silencing was achieved using commercially available ITGB1 shRNA lentiviral vector (TRCN0000029648, Sigma Aldrich).

### Lentivirus production, harvest and infection

Human umbilical vein endothelial cells (HUVECs) (C2519A) were purchased from Lonza (Australia) and cultured in endothelial growth media (EGM-2) (Lonza, CC-3162) as per manufacturers protocols. HUVECs were trypsinized using Trypsin-EDTA (Lonza, CC-5012) and maintained for up to six passages. Human embryonic kidney (HEK)-293T cells were a kind gift from Dr E. Gordon (IMB, UQ) and were maintained in Dulbecco’s Modified Eagle Medium (DMEM) (Gibco, 11995-065) containing 10% (v/v) fetal bovine serum (FBS) and 50 U/mL penicillin-streptomycin (Gibco, 15070063). HEK-293T cells were trypsinized using Trypsin-EDTA (Thermo Fischer Scientific, 15400054). All cells were cultured in 100 mm polystyrene treated cell culture dishes (Corning, CLS430167) in a humidified incubator at 37°C with 5% CO_2_.

Lentiviral constructs were packaged in HEK-293T cells by third generation lentiviral packaging plasmids, pMDLg/pRRE (Addgene, 12251), pRSV-Rev (Addgene, 12253), and pMD2.G (Addgene, 12259). Plasmids were combined with OptiMEM (Gibco, 31985-062) according to the manufacturers protocol, before being added to HEK-293T cells in DMEM. After 4 hours, HEK-293T media was replaced with DMEM supplemented with 10% (v/v) FBS. Supernatant containing lentivirus was harvested 48 hours and 72 hours post-transfection and concentrated using a Lenti-X concentrator (Takara, 631231) for 1 hour at 4°C, before being centrifuged for 45 minutes at 1500 RCF at 4°C. The supernatant was removed, before resuspension of the virus pellet in a small volume of EGM-2. Lentivirus was stored at -80°C until required.

HUVECs (P2-P3) were grown to 50% confluency for lentiviral infection. Lentivirus was thawed using pre- heated EGM-2 and added directly to HUVECs. Cells were incubated with the virus for 48 hours before the cell media was replaced with EGM-2 supplemented with puromycin (1 μg/mL, Sigma Aldrich, P8833) or neomycin (50 μg/mL, Santa Cruz, sc-29065A). After 24 hours of selection, the media was refreshed, and cells were selected for a minimum of 3 days before proceeding with experiments.

### Immunofluorescence of endothelial cell cultures

HUVECs were seeded at a density of 50,000 cells/mL and grown on untreated glass coverslips in 24-well plates until confluency was reached, with media refreshed daily. All cells were fixed at RT in 4% PFA for 10 minutes on a rotating platform and subsequently washed with PBS for a minimum of 30 minutes. Permeabilization was performed using 0.2% Triton X-100 (Sigma Aldrich, T9284) in PBS for 10 minutes before blocking using 10% goat serum (Sigma Aldrich, G6767) in PBS for 1 hour at RT. Primary antibodies in blocking buffer were incubated for 1 hour at RT, before washing with PBS for 30 minutes and incubation with corresponding secondary antibodies and anti-human CD144 conjugated antibody for 1 hour at RT. Nuclei were labelled using DAPI, incubated for 5 minutes at RT. Coverslips were mounted on glass microscope slides using Mowiol (Sigma Alrich, 81381). Antibodies and corresponding dilutions were used as described above. Confocal images were taken using a Zeiss Axiovert 200 Inverted Microscope Stand with LSM 710 Meta Confocal Scanner, using a Plan Apochromat 63X oil objective (N.A. 1.40). All imaging settings were kept constant between replicates.

### Western blotting

Cell pellets were lysed in lysis buffer containing 50 mM Tris (pH 8.0), 150 mM NaC, 1% Triton X-100, 10% glycerol and protease inhibitors cocktail (Sigma, 04693116001) or lysed directly in SDS-sample buffer containing 100 mM DTT and boiled at 95°C for 10 minutes to denature proteins. Proteins were separated by SDS_PAGE on 4-15% gradient gels (Mini-PROTEAN Precast gels, BioRad) before transfer onto a nitrocellulose membrane (Bio-Rad, 1620112) in blot buffer (48nM Tris-HCl, 39nM glycine, 0.04% SDS and 20% methanol). Blocking was completed in 5% (w/v) BSA in 0.1% Tris-based saline with Tween-20 (TBST) for 1 hour. Primary antibodies were diluted in blocking buffer and incubated overnight at 4°C followed by HRP-conjugated secondary antibodies in blocking buffer. Antibodies were removed by washing three times in 1X TBST for a minimum of 30 minutes. The HRP signals emitted were visualized by enhanced chemiluminescence (ECL, BioRad, 1705060) and imaged with a Chemidoc (BioRad). Images were analyzed using Fiji (Image J) gel analyzer software, measuring signal intensity and adjusted to the loading control (GAPDH).

### Image processing and quantification

All image processing and analysis was performed using Fiji, Image J (National Institutes of Health) and performed blinded, prior to attaining genotypes^79^.

*<u>klf2a</u>* <u>expression:</u> Images were thresholded to select for *klf2a* expression and a binary mask generated from thresholded images. Binary masks were applied to the original image before being merged with *Tg(kdrl:mCherry-CAAX)^y1^*^71^ expression to identify *klf2a* positive cells in the vasculature. Using the free- hand tool, each nucleus within the PCV was traced and saved as an ROI. Two ROIs of the highest expression within the z-stack were created per nucleus. The resulting mean gray values of these ROIs were averaged for each nucleus. The average of each embryo was quantitated by averaging *klf2a* intensity of all PCV nuclei.

*<u>Vinculinb-eGFP</u>* <u>expression:</u> A 0.5-pixel radius was used as a median filter and applied to all images. A z- projection at maximum intensity was subsequently generated for each z-stack. Vinculinb-eGFP expression was thresholded using default dark before being converted to a binary mask. Cluster count was measured within each vessel area using particle analyzer (size: 0.25 µm^2^ - 5 µm^2^).

<u>Common Cardinal Vein (CCV) EC size:</u> Using the freehand tool (Fiji, Image J), a ROI was traced around each individual cell, based on *TgBAC(ve-cad:ve-cadTS)^uq11bh^* expression at cell-cell junctions. Cell area was measured for each individual cell, and the average cell area of each embryo was reported.

<u>Lateral Dorsal Aorta (LDA) image processing:</u> Using the freehand tool, the outline of the LDA, based on *Tg(kdrl:mCherry-CAAX)^y171^* expression, was traced on each z-slice and cut out. Images were z-projected at maximum intensity for LDA vessel area and width analysis.

LDA vessel area: Using the freehand tool, a ROI was traced around the perimeter of the LDA vessel based on *Tg(kdrl:mCherry-CAAX)^y171^* expression, for a 200μm length from the base of the branch point of the vessel.

<u>LDA vessel width:</u> Vessel width was quantified using Fiji macro VasoMetrics Tool^80^, with vessel width measured at 5 μm increments. The width of the left and right branch of the LDA was recorded and the average width of both branches reported.

<u>HUVEC segmentation:</u> Images were processed through the Cellpose cell analyzer pipeline, using the cyto2 model^81,82^. The Cellpose mask outputs were then screened in ImageJ, removing any cells which were incomplete, and ROIs were updated manually if Cell Pose tracing was not accurate. For each cell, a 4 μm band was drawn inside the cell, with the banded region defined as the junctional area, and the inner area defined as the cytoplasmic area.

<u>HUVEC focal adhesion analysis:</u> Z-stack images underwent z-projection at maximum intensity. Cell traces/ROIs were selected or cells traced based on VE-cadherin cell outlines. The focal adhesion signal was thresholded using default dark before being converted to a mask. Focal adhesion count and area were measured within each cell using particle analyzer (size: 0.25 µm^2^ - infinity).

<u>Vinculin:VE-cadherin ratio:</u> A binary mask of VE-cadherin expression was generated and used to multiply by Vinculin and VE-cadherin channels. The resulting Vinculin and VE-cadherin masked images were subsequently divided to yield a Vinculin/VE-cadherin ratio, and the mean gray value was recorded.

<u>Vinculin:pPaxillin ratio:</u> A binary mask of pPaxillin expression was generated and used to multiply by Vinculin and pPaxillin individual channels. The resulting Vinculin and pPaxillin mask images were subsequently divided to yield a Vinculin/pPaxillin ratio, and the mean gray value was recorded.

<u>Nuclear KLF4 intensity:</u> DAPI expression was used to threshold for cell nuclei and analyze particles (size 10-infinity) was used to generate ROIs of all nuclei, excluding those nuclei which were not fully captured within the image area. Nuclei ROIs were then applied to the KLF4 expression and mean gray value was recorded. Analysis was performed on single z-slices.

<u>VE-cadherin intensity:</u> Images were thresholded to select for *TgBAC(ve-cad:ve-cadTS)^uq11bh^* expression and a binary mask generated from thresholded images. The binary masks were applied to the original image and a 4 μm band was drawn inside each cell based on the ROIs identified from measurement of cell size, with the banded region defined as the junctional area. Mean gray value was measured in the junctional area of each cell and recorded as the VE-cadherin intensity. VE-cadherin intensity was measured for only follower cells, with ECs of the leading edge excluded from analysis.

### Statistical analysis

All statistical analysis and graph generation was performed using GraphPad Prism9. D’Agostino-Pearson test was employed to test normal distribution of all data sets. Data sets that were normally distributed were analyzed using a One-Way ANOVA with Tukey’s multiple comparisons test where the mean of each condition was compared with the mean of every other column, or an unpaired Student’s t-test for comparison of two means. If data did not follow a normal distribution, a Mann-Whitney test was performed for comparison of two means and Kruskal-Wallis test for comparison of three or more groups. P values were represented on graphs for comparison between groups. All data is presented as Prism violin plots displaying the median and quartiles for each group. Data from independent replicates are indicated by distinct colors of the data points. For each experimental comparison, all data was acquired from sibling age-matched controls from the same clutch of embryos. HUVEC culture data was acquired from cells split from the same population and passages always matched between control and experimental groups. Image acquisition and quantification of data preceded genotyping, and analysis was completed blinded.

## Acknowledgements

Microscopy was performed in the IMB Microscopy Facility, with support from Dr Nicholas Condon for imaging and imaging analysis. We thank Prof J Vermot (Imperial College London) for kindly providing the published *klf2a:H2B-eGFP* DNA construct^62^ that we used to generate the *Tg(klf2a:H2B-eGFP)^uq5al^.* We thank The University of Queensland’s Biological Resources aquatics facility staff for zebrafish caretaking. This research was supported by a philanthropic donation from Mrs Pauline North and Mr Willian North, a Be Brave For Life Foundation micro-grant, a National Health and Medical Research Council (NHMRC) Ideas Grant (2029372), and an Australian Research Council (ARC) Discovery project (DP230100393).

