## Supplemental Figures for "Integrin β1–Talin1 at focal adhesions underpin uncontrolled endothelial cell enlargement in live cerebral cavernous malformation vasculature"

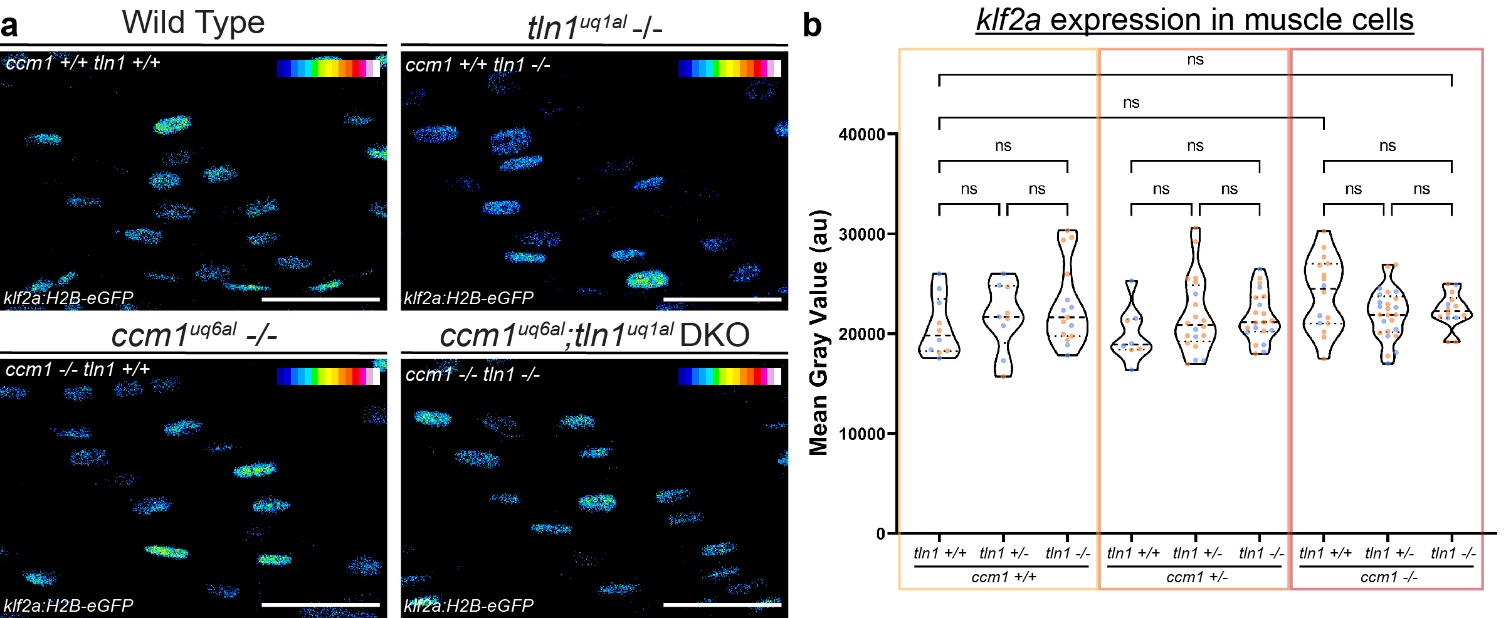

**Supplementary Figure 1: *klf2a* reporter activity in embryonic muscle. (a)** Representative images of *klf2a* expression in a single z-plane of the trunk muscle tissue in wild-type, *tln1^uq1al^ -/-*, *ccm1^uq6al^ -/-*, and *ccm1;tln1* double mutants at 48 hpf. **(b)** Quantification of *klf2a* expression in muscle cells in A; 2 biological replicates; n(*ccm1+/+;tln1+/+*) = 10, n(*ccm1+/+;tln1+/-*) = 9, n(*ccm1+/+;tln1-/-*) = 15, n(*ccm1+/-;tln1+/+*) = 8, n(*ccm1+/-;tln1+/-*) = 20, n(*ccm1+/-;tln1-/-*) = 21, n(*ccm1-/-;tln1+/+*) = 16, n(*ccm1-/-; tln1+/-*) = 25, n(*ccm1-/-;tln1-/-*) = 14; 5 nuclei per embryo. One-way ANOVA with Tukey’s multiple-comparisons test. Data represented using violin plots, small circles indicate individual data points for each replicate (color matched to replicate). ns=no significant difference.

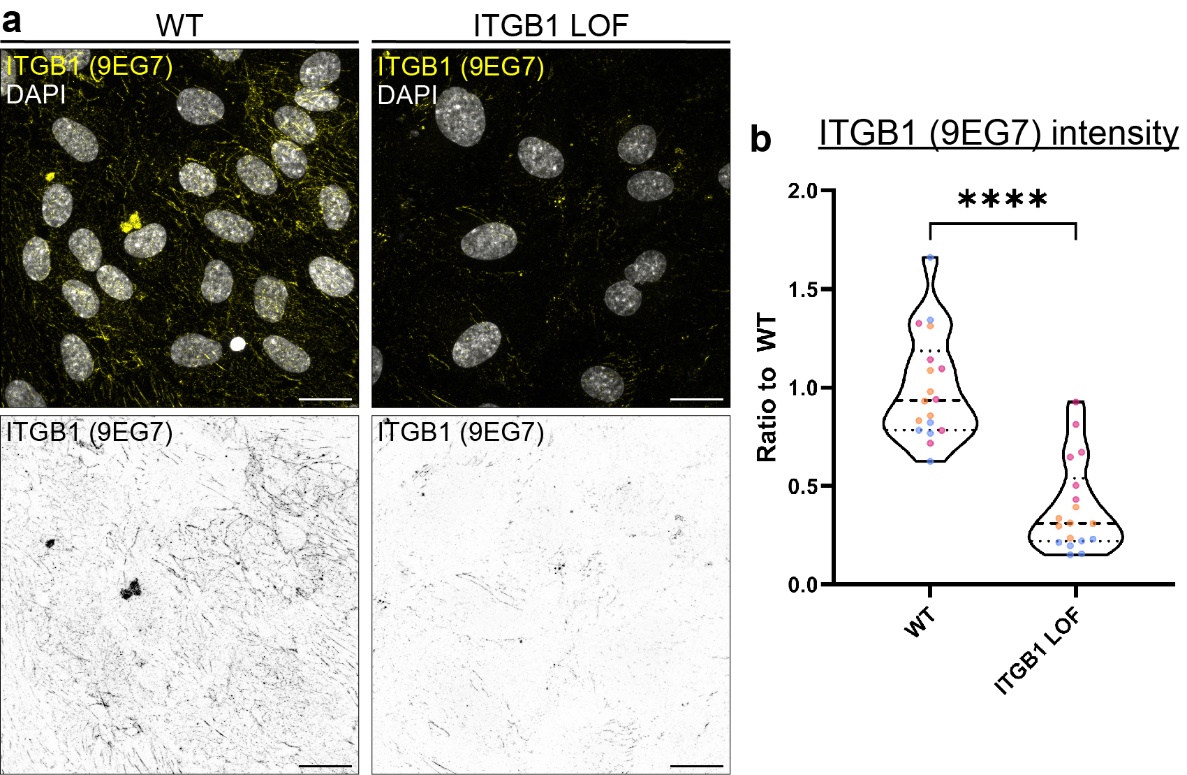

**Supplementary Figure 2: Validation of ITGB1 LOF upon shRNA lentiviral knockdown in HUVECs.** **(a)** Representative maximum z-projected confocal images of confluent monolayers of WT and ITGB1 LOF ECs, stained with the 9EG7 antibody (yellow), which specifically binds ITGB1 receptors that are in an activated confirmation, and DAPI to stain nuclei (grey) (scale bar = 20 μm). **(b)** Quantification of mean fluorescence intensity of active ITGB1 staining showing a reduction in active ITGB1 expression upon lentiviral ITGB1 shRNA infection; 3 biological replicates, 6 images per replicate. Unpaired two-tailed Student’s t-test. Data represented using violin plots, small circles indicate individual data points for each replicate (color matched to replicate). ns=no significant difference, ****p<0.0001.
